# Natural Variation in Photoprotection: Rapid NPQ Kinetics in Ferns

**DOI:** 10.1101/2025.02.07.637181

**Authors:** Nina M. Maryn, Antonio Chaparro, Audrey Short, Graham R. Fleming, Krishna K. Niyogi

**Affiliations:** Howard Hughes Medical Institute, Department of Plant and Microbial Biology, University of California, Berkeley, CA 94720; Department of Chemistry, University of California, Berkeley, CA 94720; Molecular Biophysics and Integrated Bioimaging Division, Lawrence Berkeley National Laboratory, Berkeley, CA 94720

**Author notes:** Author for correspondence: Krishna K. Niyogi. The author responsible for distribution of materials integral to the findings presented in this article in accordance with the policy described in the Instructions for Authors (https://academic.oup.com/plphys/pages/General-Instructions) is Krishna K. Niyogi.

## Abstract

Land plants perform oxygenic photosynthesis but are unable to use all of the solar radiation that they absorb on a daily basis. To minimize the production of reactive oxygen species in excess light, photosynthetic organisms use non-photochemical quenching (NPQ) mechanisms to dissipate excess excitation energy. However, the on-off transition of these mechanisms is slower than the light fluctuations themselves. In high-to-low light transitions, this can be costly to the overall productivity and carbon gain of the organism across its lifetime, because useful light energy is wasted. Here, we characterize the rapid kinetics of NPQ found in species across the fern lineage. Most of the 23 examined fern species showed faster NPQ induction and faster and more complete NPQ relaxation. Curve fitting suggested that energy-dependent quenching was the dominant contributor to the kinetics. The xerophytic fern *Astrolepis windhamii* exhibited rapid, dithiothreitol-resistant accumulation of zeaxanthin during NPQ induction, and it maintained low residual NPQ after NPQ relaxation, which however was not associated with rapid re-epoxidation of zeaxanthin. Rapid NPQ kinetics might have been an adaptive trait as ferns radiated in sunflecked forest understories during angiosperm diversification and expansion during the Cretaceous.

## Introduction

Photosynthesis is a precarious process that brings singlet excited chlorophylls into close proximity to molecular oxygen. When absorbed sunlight exceeds a plant’s capacity to process excitation energy through Photosystem II (PSII), a safety valve is required to avoid the generation of reactive oxygen species (Asada, 2006). A combination of molecular mechanisms collectively measured and referred to as non-photochemical quenching (NPQ) has evolved to dissipate this excess light energy in the PSII antenna (LHCII) as heat (Müller, Li and Niyogi, 2001; Murchie and Ruban, 2020; Cohu et al., 2013). However, the kinetics of NPQ differ between induction at the onset of high light (HL) exposure and its relaxation in low light (LL) or darkness (Müller, Li and Niyogi, 2001). In fact, there is a significant lag in the relaxation of NPQ that drives a theoretical reduction in overall photosynthetic efficiency of the organism over its lifetime, which is a limitation for crop yield (Long *et al*., 2022; Zhu et al., 2004).

Two important components of NPQ are energy-dependent (qE) and zeaxanthin-dependent (qZ) quenching, which operate on different timescales (Bassi and Dall’Osto, 2021). The rapidly reversible component, qE, is induced by excess proton buildup in the thylakoid lumen during HL, which is detected by PSII subunit S (PsbS) (Li *et al*., 2004). The more slowly reversible component, qZ, induces on a time scale of minutes and is triggered by protonation of the lumenal enzyme violaxanthin de-epoxidase (VDE) (Bratt *et al*., 1995). VDE converts violaxanthin to zeaxanthin (Zea), which has been shown to sustain and increase the magnitude of NPQ through both qE and qZ (Demmig-Adams and Adams, 1996). The re-epoxidation of zeaxanthin back to violaxanthin by zeaxanthin epoxidase (ZEP) (Schwarz *et al*., 2015), located in the stroma, is a slower reaction, which is thought to lead to slower rates of NPQ relaxation after longer periods of HL exposure that cause accumulation of Zea (Demmig-Adams and Adams, 1996).

Several approaches have been successfully applied to accelerate NPQ kinetics using the molecular components described above. Increases in biomass yield in tobacco (Kromdijk *et al*., 2016) and seed yield in soybean (De Souza *et al*., 2022) were achieved by overexpressing the Arabidopsis PsbS, VDE, and ZEP proteins to accelerate both qE and qZ. Attempts to do this in Arabidopsis (Garcia-Molina and Leister, 2020) and potato (Lehretz *et al*., 2022) accelerated NPQ recovery but did not increase yield in fluctuating light. NPQ kinetics were also accelerated by the overexpression of the Arabidopsis KEA3.2 isoform stably in Arabidopsis and transiently in *Nicotiana benthamiana* (Armbruster *et al*., 2016). Additionally, expression of PsbS from moss (Leonelli *et al*., 2016), and ZEP from the alga *Nannochloropsis oceanica* (Leonelli, Brooks and Niyogi, 2017) in *N. benthamiana* and Arabidopsis has conferred major changes in NPQ dynamics. These examples indicate that utilizing transgenic overexpression, taking advantage of natural diversity of NPQ-related genes from plant and algal species, can potentially induce a much-needed boost in crop yield by optimizing NPQ kinetics.

Here, we demonstrate that rapid NPQ kinetics, both induction and relaxation, are present in many species throughout the fern lineage. We closely examine the NPQ kinetics of the xerophytic (arid climate) fern *Astrolepis windhamii* (hereafter referred to as Astrolepis), which had the fastest NPQ kinetics of the fern species assessed. We demonstrate that rapid induction and relaxation of NPQ are likely separate mechanisms, and that the induction is linked to rapid Zea accumulation in this species upon initial high light exposure.

## Results

### In search of species with rapid NPQ

We first conducted a survey of NPQ dynamics of 77 taxonomically diverse land plant species at the University of California Botanical Garden to identify candidate species with rapid NPQ kinetics. Species were selected based on their presumed adaptation to highly fluctuating light environments, such as forest understories, rocky deserts, and dense grass stands. Chlorophyll fluorescence parameters of leaf punches were measured by imaging of pulse-amplitude modulated (PAM) fluorescence. The dark-acclimated maximum efficiency of PSII (F_v_/F_m_) of the plants ranged from 0.42 to 0.81 (Table S1). To quantify the induction and relaxation kinetics of NPQ, we plotted the ratio of the area under the curve (AUC) of normalized NPQ induction during the first 2.5 min of HL exposure (900 µmol photons m^-2^ s^-1^) to the AUC of normalized NPQ relaxation over 10 min (Fig. 1a). Species with faster NPQ induction and faster and more complete relaxation of NPQ would have a higher ratio. We were interested to determine if any particular taxonomic clades clustered in the fastest and slowest NPQ groups. 12 out of the 19 species in the upper quartile (species with the fastest NPQ) were fern species, and none in the lowest quartile were ferns.

**Fig. 1:**
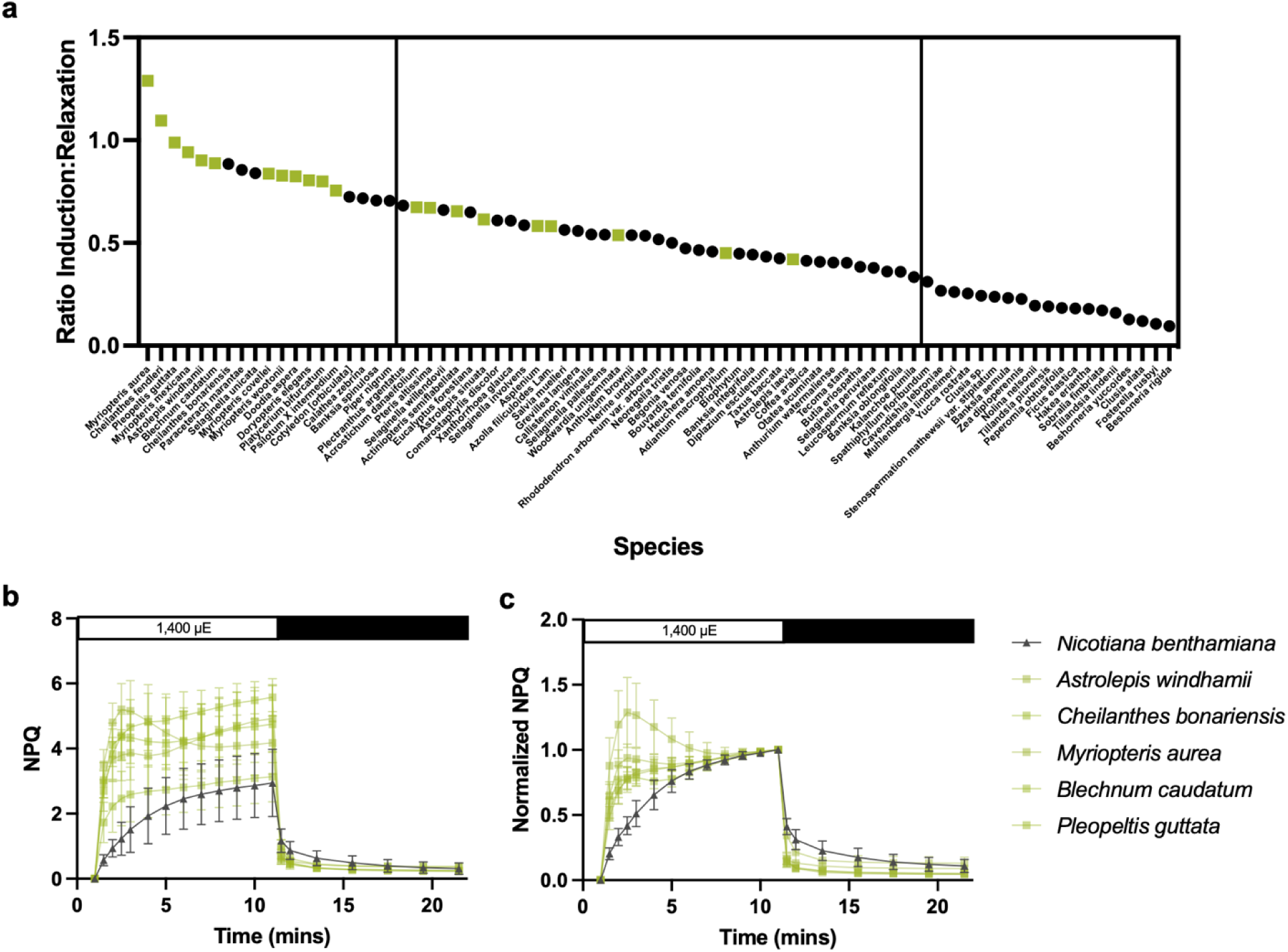
Plant NPQ kinetics survey from the UC Botanical Garden in Berkeley, CA. **a.** Ratio of area under the curve (AUC) of NPQ during induction (0-2.5 mins after 900 µmol photons m^-2^ s^-1^ HL exposure) to relaxation (10 mins of dark) for each species surveyed from the UC Botanical Garden. Upper and lower quartile are marked by vertical lines. Fern species are represented by green squares (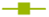). **b.** Raw NPQ for fern species in top five from the botanical garden survey, retested by pulse-amplitude modulated fluorometry (FMS2, Hansatech Instruments Ltd., Norfolk, UK) for higher resolution and compared to *Nicotiana benthamiana* tobacco (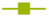 ferns, 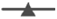 tobacco). **c.** Normalized NPQ data (normalized to the last HL timepoint) from **b**.

To confirm this phenotype with a larger sample size, we performed PAM fluorometry with whole leaf samples of the top five candidate fern species. We compared their NPQ kinetics to *N. benthamiana* grown in a greenhouse to verify that this was a reproducible, fern-specific rapid NPQ phenotype (Fig. 1b-c).

### Rapid NPQ kinetics is a common trait in the fern lineage

To determine how common rapid NPQ kinetics are across the fern lineage, we collected leaf samples from 23 species from diverse families of ferns at the University of California Botanical Garden. We chose more species from within the Pteridaceae, Eupolypod I, and Eupolypod II clades, as they are the most species rich in nature (Ranker and Haufler, 2008). F_v_/F_m_ ranged from 0.66 to 0.83 (Table S2).

All ferns examined, except for the eyelash fern *Actinopteris semiflabellata*, exhibited a more rapid NPQ induction in the first 2.5 min of HL, relative to *N. benthamiana*, which gradually reached its maximum NPQ over the course of 10 min (Fig. 2b-d). During the second HL period, ferns and *N. benthamiana* reached their maximum NPQ at a similar rate (Fig. 1b-d), which is consistent with previous reports of Zea accumulation increasing the rate of NPQ induction after HL priming in both angiosperms and fern species (Garcia-Molina and Leister, 2020; Saldaña *et al*., 2010). We fitted an exponential association curve to the first 2.5 mins of HL exposure for each species using normalized NPQ data (Table S3). The amplitude of the fit curve, A_q1_, was closely correlated to the AUC ratio measurement. *N. benthamiana* had an A_q1_ amplitude of 0.41, whereas ferns exhibited a range from 0.53 in the eyelash fern *Actinopteris semiflabellata* to 0.84 in the tree fern *Cyathea medullaris*. The majority of fern species also exhibited a lower r_q1_ than tobacco, indicating faster NPQ induction, except for *Actinopteris semiflabellata* and *Acrosticum danaeifolium*.

**Fig. 2.**
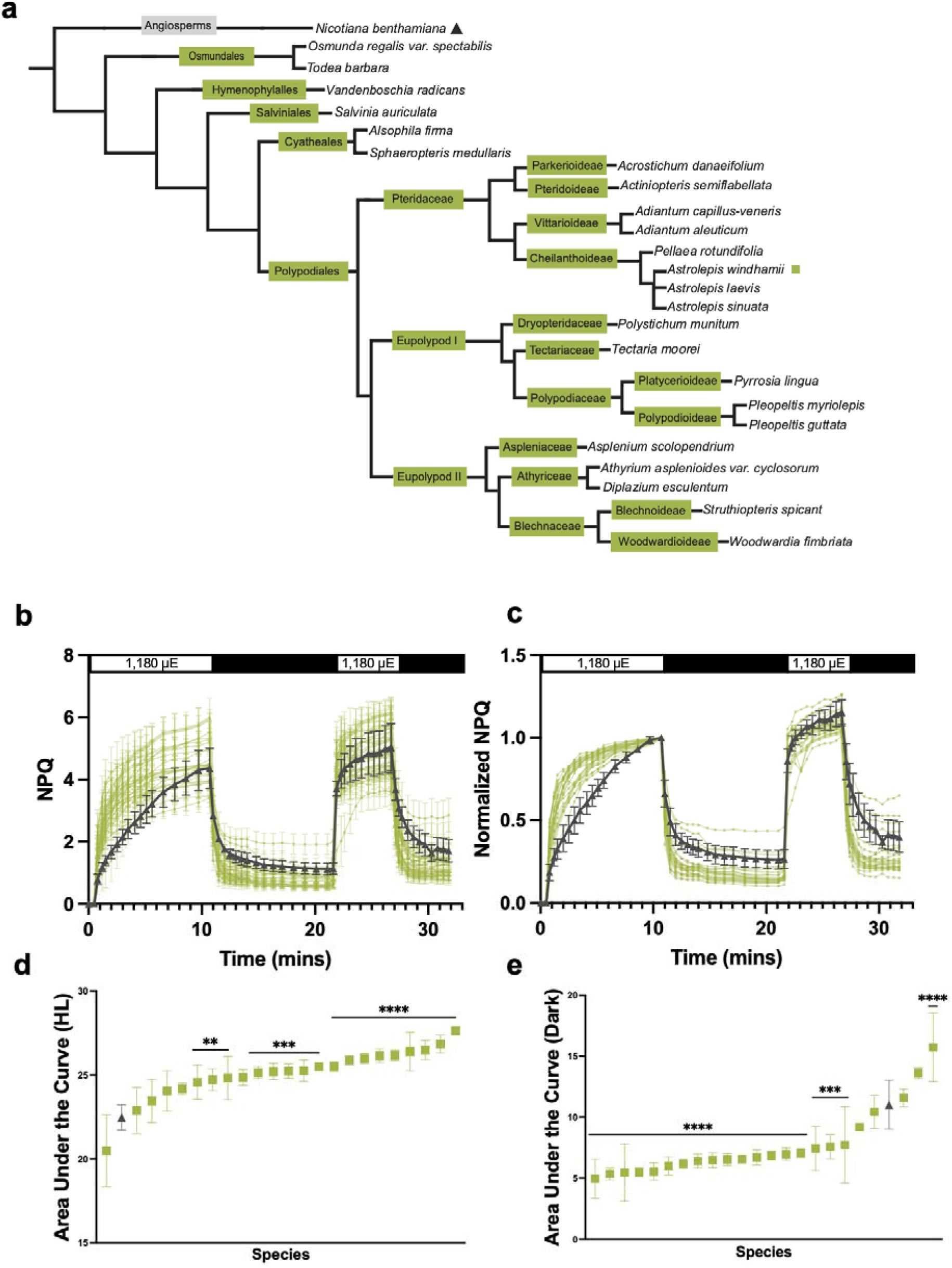
NPQ kinetics in diverse fern species. **a.** A cladogram of fern species assessed in the botanical garden survey, with *Nicotiana benthamiana* (tobacco) as an outgroup. Families and prominent subfamilies are named. *N. benthamiana* (black triangle) and *Astrolepis windhamii* (green square). **b.** NPQ kinetics (induction at 1,180 µmol photons m^-2^ s^-1^ and relaxation in the dark) of ferns and tobacco from two separate experiments. Error bars are SD (n = 3-6). **c.** Data in **b** were normalized to the end of the first HL period (10.5 mins) to show relative NPQ induction and relaxation over time. **d.** Area under the curve of normalized NPQ during HL periods for all species, ranked. Stars indicate the result of an ANOVA test, corrected for multiple comparisons to tobacco. (**** pval < 0.0001, *** pval < 0.001, ** pval < 0.01). **e.** Area under the curve of normalized NPQ during dark relaxation periods for all species, ranked. Stars are the same values as in **d**.

In both the first and second dark periods following HL exposure, 18 out of 23 fern species had significantly faster and more complete NPQ relaxation than *N. benthamiana* (Fig. 2e). This effect was more dramatic in the second dark phase, where the extent of NPQ relaxation in *N. benthamiana* was less than in the first phase, whereas ferns tended to relax back closer to their initial dark state (Fig. 2b-c). A biexponential curve was also fit to the data during the two relaxation periods in this light regime (10-20 min and 25-30 min) to approximate the contributions of qE and qZ however this method failed to adequately fit the NPQ relaxation of many of the fern species. Therefore, we compared monoexponential and biexponential fits (Table S3) and found that a monoexponential decay was a better fit for 7 out of 23 species for the first relaxation period (10-20 min in the dark). During the first relaxation period, in 21 out of 23 fern species, A_q1_ exceeded the A_q1_ of *N. benthamiana* (0.54, relative units), with 15 out of 23 fern species exceeding an A_q1_ of 0.75, indicating that over 75% of the initial relaxation of NPQ was attributable to qE rather than qZ. **τ**_q1_ ranged from 0.24-0.61 min in the ferns, whereas *N. benthamiana* fell in the middle range of these values (0.31 min), indicating similar qE kinetics. In those species that were better fit with a biexponential curve, A_q2_ ranged from as low as 0.07 in the chain fern *Woodwardia fibriata* and the maidenhair fern *Adiantum aleuticum* up to 0.40 in the slower fern *Pellaea rotundifolia*. **τ**_q2_ also had a wide range from 0.55 to 13 mins. This suggests that qZ can be a significant contributor to fern NPQ relaxation, but this varies widely by species. A_q3_, the contribution of slowly reversible photoinhibitory quenching (qI), was lower than that of tobacco (0.26) in all but two fern species, the aquatic fern *Salvinia auriculata* and the maidenhair fern *Adiantum aleuticum*. In the second dark relaxation period, 18 out of 23 fern species and *N. benthamiana* were better fit with the monoexponential curve (Table S4), suggesting that qE and qI were the dominant contributors to the relaxation kinetics after a previous exposure to fluctuating light.

### Characterizing NPQ kinetics in *Astrolepis windhamii*

We next conducted an in-depth investigation of the NPQ kinetics of a single fern species to delve into the mechanics of fern NPQ. We chose *Astrolepis windhamii* (Astrolepis) because it (1) exhibited the fastest relaxation phenotype of all ferns assessed in our botanical garden survey and (2) had young, similarly aged fronds to be able to control for age and growth conditions at the times of sampling. We assessed Astrolepis NPQ in four light regimes, and greenhouse-grown *N. benthamiana* was subjected to the same regimes for comparison. In all experiments, there was no significant difference between the F_v_/F_m_ values for Astrolepis and *N. benthamiana*, which were all above 0.8 on average (Fig. S1a-d). The fluorescence emission for Astrolepis was also much higher than for *N. benthamiana*, but this is likely accounted for by leaf thickness with higher chlorophyll per leaf area in Astrolepis (Fig. S1e-h).

The light response of NPQ was measured at 15 light intensities between 5 µmol photons m^-2^ s^-1^ and 2,200 µmol photons m^-2^ s^-1^. NPQ was found to be significantly higher (p < 0.0001) in Astrolepis at all light intensities measured above 60 µmol photons m^-2^ s^-1^ (Fig. 3a), which was accompanied by a significantly lower (p < 0.0001) operating efficiency of PSII (**Φ**PSII) (Fig. 3b).

**Fig. 3.**
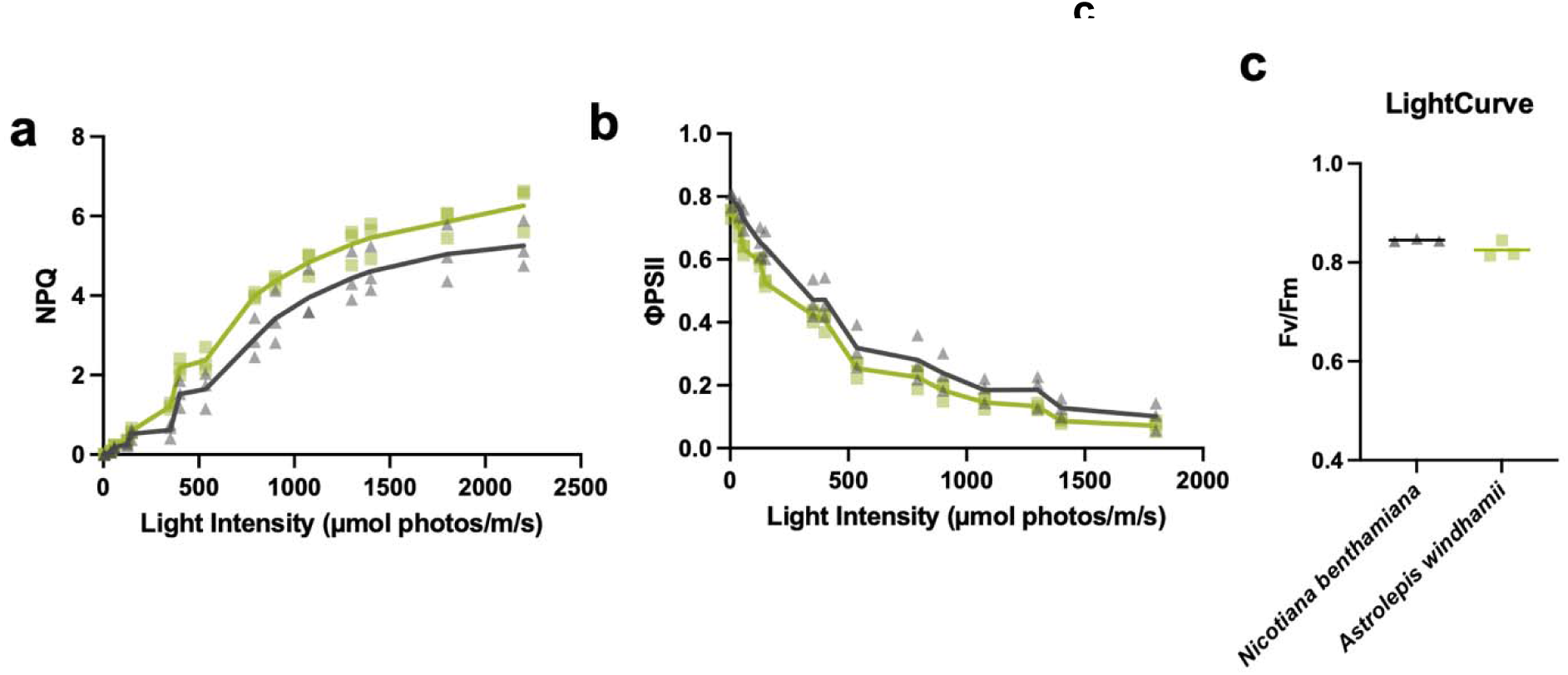
NPQ and φPSII at increasing light intensities in Astrolepis. **a.** Light response curve of NPQ values after 5 min of acclimation at increasing light intensities in tobacco and Astrolepis (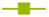 Astrolepis, 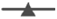 tobacco) **b.** φPSII values from the light response experiment in **a**. **c.** Corresponding Fv/Fm values for Nicotiana and Astrolepis.

We examined NPQ during a 10 min HL (1,180µmol photons m^-2^ s^-1^)-dark cycle followed by a 5 min HL-dark cycle and obtained results that supported our initial measurements using video imaging of chlorophyll fluorescence (Fig. 4a). This was accompanied by a slightly higher **Φ**PSII measured during the dark relaxation periods in Astrolepis compared to *N. benthamiana*, but no difference in HL (Fig. 2b). The induction of NPQ during the first HL exposure was faster in Astrolepis compared to *N. benthamiana*, with the AUC in the first 2 min of HL reaching 9.2 in Astrolepis on average, compared to just 3.9 in *N. benthamiana* (Fig. 4c). Similarly, the residual NPQ (as measured by cumulative normalized NPQ AUC in the dark) was 2.2 times higher in *N. benthamiana* compared to Astrolepis, indicating a much faster and more complete NPQ relaxation in Astrolepis (Fig. 4d).

**Fig. 4.**
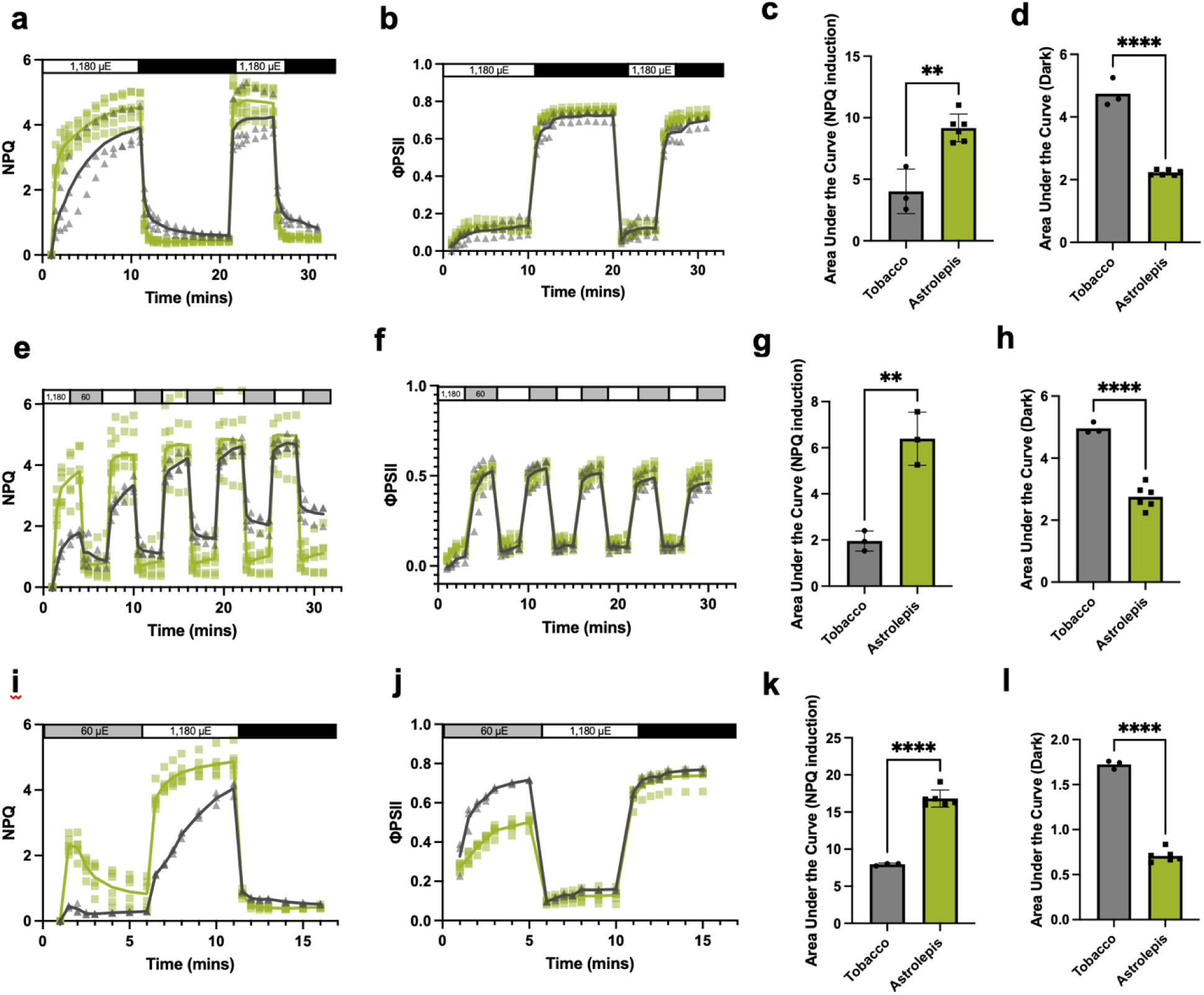
Characterization of NPQ kinetics in Astrolepis. **a**. NPQ kinetics in Astrolepis and tobacco (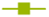 Astrolepis, 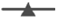 tobacco) over a 10 min HL - 10 min dark period followed by 5 min HL - 5 min dark period of 1,180 µmol photons m^-2^ s^-1^. **b.** φPSII values from the experiment in **a**. **c.** Area under the curve during NPQ induction (first 2.5 min of HL) in Astrolepis versus tobacco from data in **a**. **d.** Area under the curve during normalized NPQ relaxation (NPQ in the dark) in Astrolepis versus tobacco from data in **a**, normalized to the last HL timepoint. **e.** NPQ kinetics in Astrolepis and tobacco over five 2.5 HL – 2.5 LL periods of 1,180 µmol photons m^-2^ s^-1^ and 60 µmol photons m^-2^ s^-1^, respectively. **f.** φPSII values from the experiment in **e**. **g.** Area under the curve during NPQ induction (first 2 min of HL) in Astrolepis versus tobacco from data in **e**. **h.** Area under the curve during NPQ relaxation (cumulative NPQ in LL) in Astrolepis versus tobacco from data in **e**, normalized to the last HL timepoint. **i.** NPQ kinetics in Astrolepis and tobacco over a regime of 5 mins of LL (60 µmol photons m^-2^ s^-1^), 5 min of HL (1,180 µmol photons m^-2^ s^-1^) and 5 min of dark. **j.** φPSII values from the experiment in **i**. **k.** Area under the curve during NPQ induction (first 2 min of HL) in Astrolepis versus tobacco from data in **i**. **k.** Area under the curve during normalized NPQ relaxation (NPQ in the dark) in Astrolepis versus tobacco from data in **i**, normalized to the last HL timepoint.

Interestingly, in a more rapid 2.5 min HL-LL (60 µmol photons m^-2^ s^-1^) cycling regime, Astrolepis was able to maintain an almost square-wave NPQ kinetics phenotype over five cycles, achieving its maximum NPQ almost immediately and returning to its minimum NPQ. In contrast, *N. benthamiana* exhibited a gradual increase in maximum and minimum NPQ over the course of many cycles (Fig. 4e). In this experiment, **Φ**PSII was indistinguishable between the species (Fig. 4f). Over several cycles, the total residual NPQ was significantly lower as measured by cumulative AUC and total NPQ in HL was higher for Astrolepis (Fig 4g-h), as we saw in longer light periods. These NPQ phenotypes in longer HL periods and fluctuating light regimes were confirmed by time-correlated single photon counting (TCSPC) measurements of chlorophyll fluorescence lifetimes (Fig. S2), indicating that the phenotypes are consistent across different types of measurements and not due to rapid chloroplast movements, which are ruled out via TCSPC.

Finally, we wanted to see how LL to HL transitions affect fern NPQ relaxation in the dark. Surprisingly, we consistently found a strong rapid NPQ induction of around 2.2 upon transition from dark to LL in Astrolepis, whereas *N. benthamiana* had a transient NPQ in LL of only 0.4 (Fig. 4i), which is similar to, but more robust than what has been observed in Arabidopsis (Finazzi *et al*., 2004; Kalituho, Beran and Jahns, 2007). This transient NPQ slowly relaxed during the LL period of 5 min and did not inhibit full NPQ induction during a subsequent HL period in Astrolepis (Fig 4k), though it did seem to correlate with a slower induction in *N. benthamiana* (Fig. 4i). However, there was a significantly lower (p < 0.0001) **Φ**PSII during this LL period in Astrolepis (Fig. 4j). Normalized relaxation in Astrolepis during the final dark exposure was still significantly faster (Fig 4l), but in this case the relaxation was attributable to the higher NPQ of Astrolepis in HL than *N. benthamiana*, with both having similar residual NPQ levels in the dark (Fig 4i).

### Astrolepis NPQ induction is correlated to rapid, DTT-resistant zeaxanthin accumulation

To examine whether rapid xanthophyll cycling could be a factor in fern NPQ kinetics, we performed high-performance liquid chromatography (HPLC) analysis of photosynthetic pigments in Astrolepis. Figure 5a shows the results of a time course of Astrolepis and *N. benthamiana* leaf discs exposed to LL (60 µmol photons m^-2^ s^-1^) or HL (∼1,100 µmol photons m^-2^ s^-1^). After an overnight dark acclimation, neither Astrolepis and *N. benthamiana* had Zea or antheraxanthin present. However, after just 1.5 min of HL exposure, Astrolepis had reached a DES of 0.18, while *N. benthamiana* still had a DES of 0. This DES in Astrolepis was maintained up to 10 min in HL, while *N. benthamiana* reached a mean DES of 0.03 after 10 min. During a subsequent dark period, no re-epoxidation of Zea to Vio occurred within 1.5 min in either species. After 10 min of HL followed by a 10 min dark period and a subsequent 1.5 min of HL, Astrolepis reached a DES of 0.26. *N. benthamiana* in this condition only reached a DES of 0.09. Additionally, dark-acclimated Astrolepis was able to achieve a DES of 0.15 in LL after only 1.5 min, suggesting that Zea may contribute to the transient NPQ observed in LL (Fig. 4i). The concentrations of lutein, neoxanthin, and chlorophyll *b* (relative to chlorophyll *a*), were higher in Astrolepis than *N. benthamiana* (Fig. 5b), which indicates potentially a larger PSII antenna size in Astrolepis, harboring these pigments.

**Fig 5.**
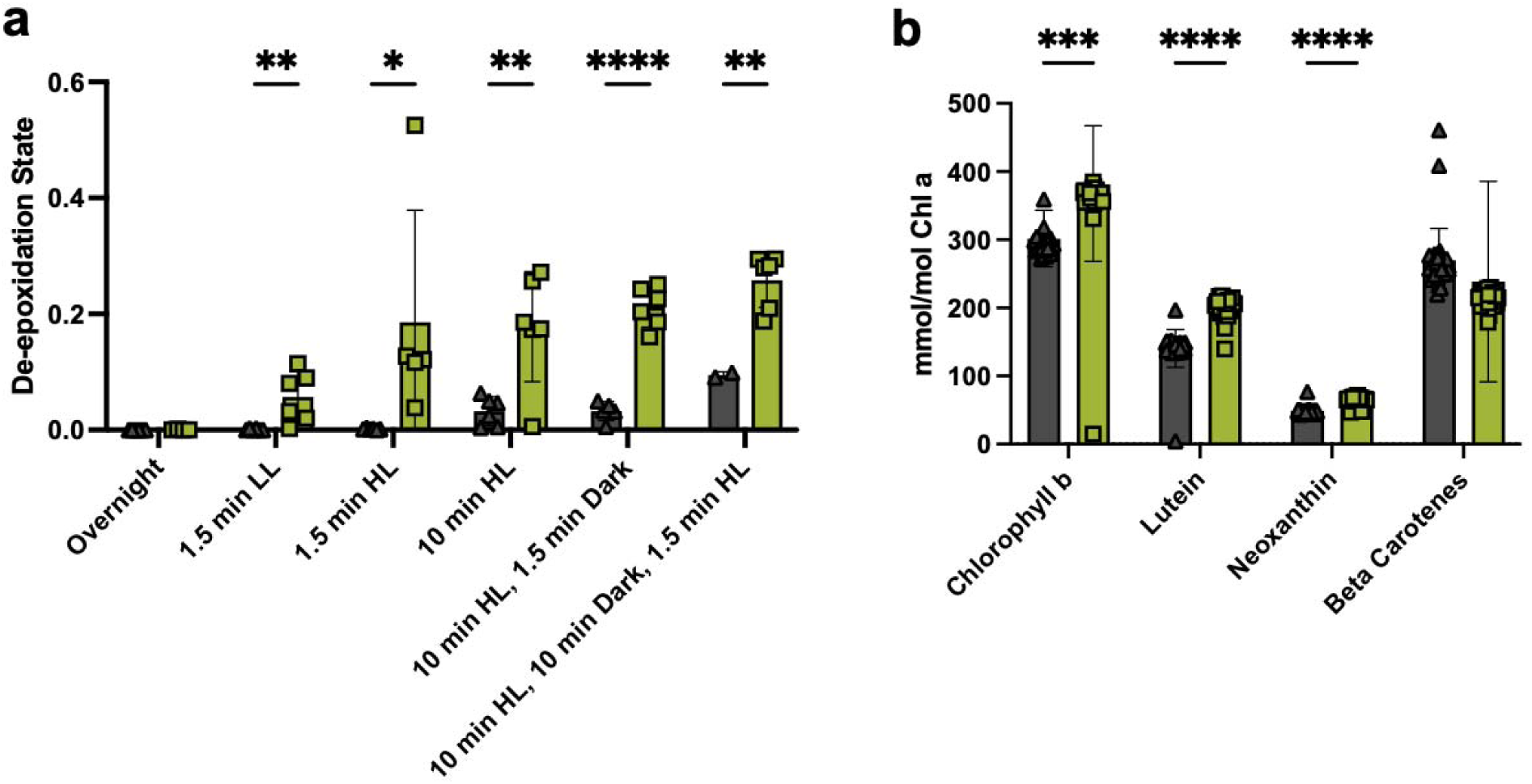
Pigment analysis of tobacco and Astrolepis. **a.** De-epoxidation state (--------------------) in tobacco and Astrolepis after overnight dark acclimation, followed by exposure to LL (60 µmol photons m^-2^ s^-1^) or HL (1,180 µmol photons m^-2^ s^-1^) or fluctuating light. **b.** Chlorophyll *b* and non-xanthophyll-cycle carotenoids relative to chlorophyll *a*, from the samples in **a**.

To test whether NPQ in Astrolepis is more reliant on qE or qZ, we treated leaf discs with three inhibitors and measured NPQ kinetics during 5 min LL, followed by two cycles of 5 min HL and 5 min dark. We first used the PSII inhibitor DCMU (1 mM) to demonstrate that we could successfully infiltrate the leaves of Astrolepis, given that we were concerned about cell wall thickness and the ability to infiltrate the chloroplast (Fig. 6a). DCMU abolished nearly all NPQ in both Astrolepis and *N. benthamiana*, including the transient NPQ in LL in Astrolepis. We then tested the uncoupler nigericin (10 µM), which should prevent the buildup of the proton gradient across the thylakoid membrane. The transient NPQ in LL was unaffected by nigericin at this concentration, but NPQ in HL was reduced by about 50% in both species, with no effect on the relaxation kinetics (Fig. 6b).

**Fig 6.**
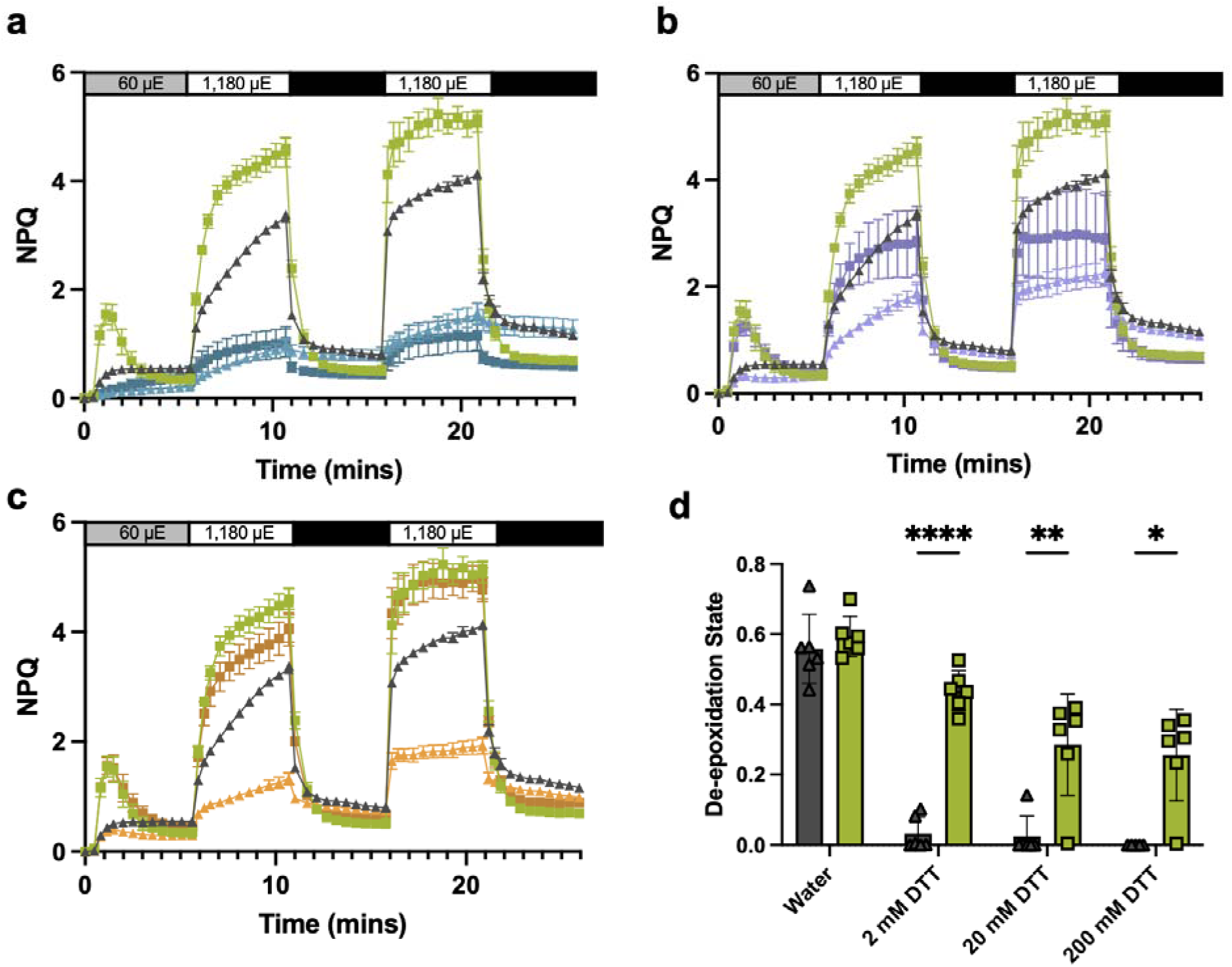
Chemical inhibition of NPQ in tobacco and Astrolepis. **a-c.** NPQ kinetics during 5 min LL, followed by two cycles of 5 min HL and 5 min dark. NPQ in tobacco and Astrolepis infiltrated with either 150 mM sorbitol (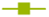 Astrolepis, 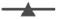 tobacco) or the following chemical inhibitors. **a.** Tobacco and Astrolepis infiltrated with 1 mM DCMU (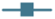 Astrolepis, 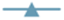 tobacco). **b.** Tobacco and Astrolepis infiltrated with 10 µM nigericin (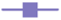 Astrolepis, 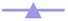 tobacco). **c.** Tobacco and Astrolepis infiltrated with 2 mM DTT (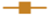 Astrolepis, 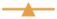 tobacco). **d.** De-epoxidation state derived from HPLC analysis of tobacco and Astrolepis leaf discs infiltrated with water, 2 mM DTT, 20 mM DTT or 200 mM DTT after 1 hour of HL exposure.

Finally, the VDE inhibitor DTT (2 mM) was used to examine whether Astrolepis NPQ depends on its rapid Zea accumulation. Surprisingly, we found that Astrolepis NPQ was not significantly hindered in the presence of DTT (Fig. 6c). To test whether DTT was able to inhibit Astrolepis VDE, we exposed leaf discs to 2 mM, 20 mM or 200 mM DTT (or water as a control) and then to 1 hour of HL (1,100 µmol photons m^-2^ s^-1^). HPLC analysis showed that the DES of controls for both Astrolepis and tobacco was approximately 0.5-0.6. De-epoxidation of Vio to Zea was abolished entirely in tobacco at all DTT concentrations, whereas Astrolepis maintained a DES of 0.44 in 2 mM DTT (Fig. 6d), suggesting that the lack of inhibition of NPQ by DTT was due to the inability of DTT to inhibit VDE in Astrolepis. Even at 20 mM and 200 mM DTT, the DES of Astrolepis was only reduced to ∼0.27 (Fig. 6d).

## Discussion

Photosynthetic energy conversion is a major limiting step in biomass accumulation in crops. In most plants, less than 1% of the energy of incident sunlight is converted into chemical energy in the form of plant biomass, and typical field conditions result in light saturation of photosynthesis at around 25% of the maximum solar flux (Jansson *et al*., 2010). The excess absorbed light energy is dissipated as heat as a result of NPQ, however the relatively slow turning off of NPQ under low light conditions is modeled to limit CO_2_ assimilation by 30% in field conditions with fluctuating light (Zhu *et al*., 2004). Therefore, understanding how to more optimally tune NPQ in crops to available light levels is a burgeoning area of research with high potential yield improvements. Recent studies have shown that accelerating the relaxation kinetics of NPQ can improve the efficiency of photosynthesis in fluctuating light (Kromdijk *et al*., 2016; De Souza *et al*., 2022), at least in some species, thus identifying NPQ kinetics as a novel trait for crop improvement. This work raises questions about how much natural variation exists in NPQ kinetics of plants, both within individual crop species and across different plant lineages. Here, we identify ferns as a group of plants with exceptionally rapid NPQ kinetics.

Our results show that rapid NPQ kinetics is a broadly conserved trait among ferns (Fig. 1), suggesting that it has a genetic basis. Ferns may have evolved this trait and adapted to fluctuating light during the evolution of extant fern lineages in the understory of angiosperm forests during the Cretaceous period (Schneider et al., 2004; Schuettpelz and Pryer, 2009). Subsequent diversification and radiation of fern species into different habitats could have occurred with the retention of rapid NPQ kinetics in most lineages. Ferns are well-documented to have strong limitations on rates of carbon assimilation, due in large part to thick mesophyll cell walls that inhibit gas exchange as an adaptation for drought resilience (Tosens *et al*., 2016). In response, different lineages have adapted their light reactions to optimize the light absorption to limits on carbon fixation in the Calvin-Benson cycle (Gago *et al*., 2019). Fern species have also been shown to differ in electron transport rate and photosystem II efficiency, using differing strategies to compete for shade or sun niches in the forest understory (Saldaña *et al*., 2010).

The mechanism of the rapid NPQ kinetics in Astrolepis appears to be different from transgenic “VPZ” plants that overexpress VDE, PsbS, and ZEP proteins. The relaxation kinetics in ferns appear to be mono-exponential in most cases, with a residual NPQ much lower than observed in VPZ plants (Kromdijk *et al*., 2016; De Souza *et al*., 2022), and re-epoxidation of Zea is not occurring on this timescale in Astrolepis (Fig. 5A). This indicates that the rapid NPQ relaxation kinetics is related to the Zea and PsbS-dependent qE component of NPQ, which may be a higher proportion (if not all) of the Zea-dependent mechanisms in these fast NPQ ferns, rather than the more slowly reversible Zea-dependent component, qZ. Induction of NPQ is highly correlated to the de-epoxidation state in Astrolepis, suggesting that NPQ capacity is related to the proportion of the VAZ pool that is comprised of Zea, as is well established in the literature (Adams, *et al*., 1999). Further elucidation of the mechanism of rapid NPQ kinetics in ferns could provide new strategies for engineering this trait in crop plants.

Other studies of photosynthesis and photoprotection in different plant species have revealed substantial natural variation in NPQ amplitude and kinetics, especially in the qZ component of NPQ (Johnson et al., 1993; Sello et al., 2019; Demmig-Adams et al., 2020), suggesting that interspecies variation is explained in part by life cycle and environmental stress adaptations. Forest understory plants that experience sunflecks tend to retain Zea after the first sunfleck of the day, and they exhibit rapid qE induction to provide photoprotection and rapid qE relaxation to maintain photosynthetic efficiency in the shade (Watling et al., 1997; Logan et al., 1997; Adams et al., 1999; Demmig-Adams et al., 2020). In some cases, however, Zea retention is associated with decreases in PSII efficiency following sunflecks (Watling et al., 1997; Adams et al., 1999), which is likely related to slowly reversible qZ. In terms of life cycle, annuals tend to thrive in high light and exhibit lower shade tolerance and lower photoprotective capacity, which is correlated with lower levels of Zea accumulation and higher photosynthetic energy conversion used in carbon fixation (Demmig-Adams and Adams, 1992). Evergreens, by contrast, generally have higher photoprotective capacity and slower carbon assimilation rates but can readily acclimate to high light or shade and optimize their growth under favorable conditions (Verhoeven, 2014). Previous studies of *Monstera deliciosa*, an understory plant, have found retained Zea in the dark and sustained levels of high NPQ after high to low light or dark transitions (Demmig-Adams *et al*., 2006). Fern species are either deciduous or evergreen, with most exhibiting an evergreen lifestyle. While we do see retention of high Zea accumulation after dark relaxation in Astrolepis, evergreen ferns including Astrolepis seem to not have evolved a propensity toward sustained NPQ in dark transitions, which could be related to the apparent low amplitude of qZ. The commonality of rapid NPQ kinetics and low residual NPQ in low light and darkness across the fern lineage, regardless of life cycle adaptation, seems to indicate a de-coupling of Zea retention and relaxation kinetics driven by a common genetic origin.

Compared to examination of interspecies variation, intraspecies analysis of NPQ kinetics offers several advantages in identifying the underlying molecular mechanisms. For example, intraspecies studies have demonstrated that there is an inherent flexibility within the individual organism as it acclimates to its environment. For example, in Arabidopsis, VAZ pool size is regulated by long-term acclimation of an individual to high light (Kawabata and Takeda, 2014), affecting NPQ capacity as well as residual NPQ in low light, and acclimation to high or low temperature and day length during development can regulate leaf architecture in a way that changes the abundance of mesophyll cells, altering the amount of light exposure the photosystems will experience in various layers of the leaf or canopy (Cohu et al., 2013). While these relatively dramatic phenotypic acclimations can occur within an individual ecotype, variation of NPQ kinetics within species appears to be relatively low compared to what we have found in ferns. Genome-wide association studies and quantitative trait analyses have revealed that differences in NPQ kinetics within plant species can be traced mainly to regulation of PsbS expression (Umate, 2010). Studies of soybean and maize crossbred accession panels showed relatively little variation in NPQ kinetics, particularly in residual NPQ in dark or low light, implying that these crops may not possess enough genetic variation between accessions to use breeding as an effective method to optimize NPQ kinetics (Wang *et al*., 2020; Ferguson *et al*., 2023), at least to the same extent as that achieved in engineered VPZ plants.

To further investigate the mechanism of rapid NPQ kinetics in ferns, it will be critical to examine the activity of important molecular components of NPQ in ferns compared to angiosperms. PsbS and VDE are the canonical major players in qE induction, so examining whether these proteins have evolved to act more quickly in ferns in response to ΔpH, for example, would be a logical next step. Similarly, the light-harvesting complexes in the PSII antenna are the site of NPQ, and these proteins may have evolved in ferns to rapidly undergo conformational changes required for qE and rely less on qZ for photoprotection. Understanding the mechanisms of rapid NPQ kinetics in ferns may also help us to better understand the complex interactions of these molecular players in higher plants and to genetically engineer these mechanisms in crops to optimize their photosynthesis in fluctuating light. Alternatively, the rapid NPQ relaxation kinetics and low residual NPQ in ferns could turn out to be a highly complex mechanism emerging from the context of the fern thylakoid membrane, which could lead to new understanding of how light harvesting is regulated in plants.

## Methods

### Plant collections

Whole leaves were harvested from plants and kept in a plastic bag with a moist paper towel in the dark until they could be brought back to the lab for analysis or overnight dark acclimation. Diverse fern species were chosen based on taxonomic diversity from the PteridoPortal database (https://pteridoportal.org/portal/index.php) and presence/abundance of plant material at the University of California Botanical Gardens in Berkeley, California (https://botanicalgarden.berkeley.edu/collections). A complete list of species and accession numbers from the UC Botanical Garden can be found in Extended Data 1.

### NPQ kinetics measurements

NPQ measurements were conducted for the species’ NPQ surveys, as well as tobacco transient expression, were conducted on hole punches from detached leaves floated on tap water in a 96 well plate. For the original species survey, Samples were dark acclimated for 30 mins prior to measurement on the IMAGING-PAM M-Series (Heinz Walz, Effeltrich,Germany) using 10 min of blue actinic light (900 µmol photons m^2^ sec^-1^), then 10 min of dark recovery. For the fern species survey and subsequent measurements for transient expression experiments in tobacco, samples were dark acclimated for 40 mins prior to measurement on the IMAGING-PAM M-Series (Heinz Walz, Effeltrich,Germany) using 10 min of blue actinic light (1,100 µmol photons m^2^ sec^-1^), then 10 min of dark recovery, followed by another cycle of 5 mins each. Photosynthetic parameters were calculated as described in Brooks and Niyogi (2011).

Tobacco and Astrolepis NPQ measurements were conducted on detached leaves using a Hansatech FMS-1 fluorometer (Hansatech Instruments Ltd., Norfolk, UK). Leaves were hydrated and dark acclimated for at least 1 hour prior to measurements. High light measurements were conducted at 1,100 µmol photons m^2^ sec^-1^ and low light measurements at 60 µmol photons m^2^ sec^-1^ unless otherwise stated.

### Time-correlated single photon counting

Time-correlated single photon counting (TCSPC) was used on whole fern leaves, which were kept moistened and in the dark, to collect chlorophyll fluorescence lifetime snapshot measurements. TCSPC while similar to PAM is not sensitive to changes in absorbance from chloroplast avoidance or changes in fluorescence yield. The technique used here is similar to Steen *et al*. (2020). Briefly, TCSPC results in a histogram of Chl *a* fluorescence decay, which is then fit to a biexponential decay function yielding a lifetime (τ_avg_). These fluorescence lifetimes were captured at 15 second intervals for 25 minute long experimental runs, resulting in snapshots of fluorescence trajectory that track the changes in the fluorescence lifetime as a function of HL exposure. The amplitude-weighted average lifetime of the Chl *a* fluorescence decay is converted into a unitless form, similar to that measured in the conventional pulse-amplitude modulation technique using the following equation: 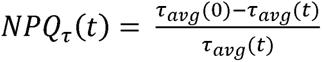 where τ_avg_(0) and τ_avg_(t) are the average lifetimes in the dark and at any time point *t* during the HL exposure, respectively (Sylak-Glassman *et al*., 2016).

An ultrafast Ti:sapphire coherent Mira 900f oscillator was pumped using a diode laser (Coherent Verdi G10, 532 nm). The center wavelength of the oscillator was 808 nm with a full width at half maximum of 9 nm. After frequency doubling the wavelength to 404 nm with a β-barium borate crystal, the beam was split between a sync photodiode, which was used as a reference for snapshot measurements, and the sample. To control exposure of the sample to the actinic light, three synchronized shutters located in the laser path, actinic light path, and path between the sample and the microchannel plate-photomultiplier tube detector (Hamamatsu R3809U) were controlled by a LABVIEW software sequence. The detector was set to 680 nm to measure Chl *a* emission. During each snapshot, the laser and detection shutters were opened, allowing an excitation pulse with a power of 1.7 mW to saturate the reaction center for 1 second while the emission was recorded (Schansker, Tóth and Strasser, 2006). During HL periods, samples were exposed to white light with an intensity of 1000 μmol photons m^-2^ s^-1^ (Leica KL 1500 LCD, peak 648 nm, FWHM 220 nm) by opening the actinic light shutter.

### Pigment analysis

Leaf discs (0.5 cm^2^) were frozen in liquid nitrogen and beat in FastPrep Lysing matrix D (tobacco) or Lysing matrix J (Astrolepis) (MP Biomedical). 150 µl of ice cold acetone was added to each sample and bead as described above. Cell debris was pelleted at 14,000 rpm for 5 min and the supernatant was collected. Samples were extracted a second time as described above, and supernatants were pooled to total 300 µl. Samples were analyzed with a Spherisorb S5 ODS1 4.6- × 250-mm cartridge column (Waters, Milford, MA) at 28°C as previously described (Müller-Moulé, Conklin and Niyogi, 2002).

### Chemical inhibitors

NPQ inhibition experiments were performed by perforating the bottom of Astrolepis leaves (which are scaly and hydrophobic) with a dermabrasion roller prior to hole punching in order to infiltrate the very thick leaves. This was compared to tobacco which were simply hole punched directly into the chemical solutions. The DTT experiments were performed at 200 mM, 20 mM and 2 mM concentrations, DCMU at 1 mM concentration, and nigericin at 10 µM and 50 µM, all diluted in 150 mM sorbitol in water was used to infiltrate the control leaf discs. Leaf discs were incubated in solution for 40 min in the dark prior to being placed on solution-wetted filter paper for NPQ measurements and subsequent freezing for pigment analysis.

## Acknowledgments

We thank Holly Ward and Clare Loughran for providing access to plants at the University of California Botanical Garden. This work was supported by a subaward from the University of Illinois as part of the research project Realizing Increased Photosynthetic Efficiency (RIPE), funded from 2017-2023 under grant number OPP1172157, by the Bill & Melinda Gates Foundation, Foundation for Food and Agriculture Research, and the U.K. Government’s Department for International Development, and by the project Realizing Increased Photosynthetic Efficiency (RIPE), that is funded by Bill & Melinda Gates Agricultural Innovations grant investment 57248, awarded to the University of California, Berkeley by the University of Illinois, USA. The TCSPC experiments were supported by the U.S. Department of Energy, Office of Science, through the Photosynthetic Systems program in the Office of Basic Energy Sciences. This material is based upon work by N.M.M. supported by the National Science Foundation Graduate Research Fellowship Program under Grant No. DGE 2146752. Any opinions, findings, and conclusions or recommendations expressed in this material are those of the author(s) and do not necessarily reflect the views of the National Science Foundation. K.K.N. is an investigator of the Howard Hughes Medical Institute. This article is subject to HHMI’s Open Access to Publications policy. HHMI lab heads have previously granted a nonexclusive CC BY 4.0 license to the public and a sublicensable license to HHMI in their research articles. Pursuant to those licenses, the author-accepted manuscript of this article can be made freely available under a CC BY 4.0 license immediately upon publication.

## Author contributions

N.M.M. and K.K.N. designed the research; N.M.M., A.C., and A.S. performed research; N.M.M. and A.S. analyzed data; N.M.M. wrote the paper with input and edits from A.C., A.S., G.R.F., and K.K.N.

---

**Table S1:** F_v_/F_m_ and n for each species sampled for plant NPQ survey.

| Species | Mean $F_v/F_m$ | SD | $n$ |
| --- | --- | --- | --- |
| *Astrolepis laevis* | 0.42 | 0.02 | 3 |
| *Fosterella rusbyi* | 0.52 | 0.03 | 3 |
| *Zea diploperennis* | 0.53 | 0.12 | 3 |
| *Doodia aspera* | 0.55 | 0.04 | 3 |
| *Muhlenbergia lindheimeri* | 0.56 | 0.15 | 3 |
| *Ficus elastica* | 0.57 | 0.05 | 3 |
| *Bouvardia ternifolia* | 0.58 | 0.06 | 3 |
| *Hakea eriantha* | 0.59 | 0.04 | 3 |
| *Tecoma stans* | 0.59 | 0.06 | 3 |
| *Beshoneria rigida* | 0.59 | 0.06 | 3 |
| *Banksia aemula* | 0.61 | 0.04 | 3 |
| *Banksia oblongifolia* | 0.62 | 0.07 | 3 |
| *Tillandsia piurensis* | 0.64 | 0.07 | 3 |
| *Myriopteris woottonii* | 0.65 | 0.02 | 3 |
| *Astrolepis sinuata* | 0.65 | 0.07 | 3 |
| *Taxus baccata* | 0.65 | 0.08 | 3 |
| *Clusia alata* | 0.65 | 0.03 | 3 |
| *Neoregelia tristis* | 0.66 | 0.07 | 6 |
| *Azolla filiculoides* Lam. | 0.67 | 0.17 | 3 |
| *Myriopteris aurea* | 0.68 | 0.02 | 3 |
| *Myriopteris covillei* | 0.68 | 0.01 | 3 |
| *Woodwardia unigemmata* | 0.68 | 0.01 | 3 |
| *Beshomeria yuccoides* | 0.68 | 0.01 | 3 |
| *Leucospermum reflexum* | 0.69 | 0.08 | 3 |
| *Myriopteris mexicana* | 0.70 | 0.03 | 3 |
| *Banksia spinulosa* | 0.70 | 0.07 | 6 |
| *Tillandsia lindenii* | 0.70 | 0.03 | 3 |
| *Stenospermatum mathewsii* var. *stipitatum* | 0.70 | 0.05 | 3 |
| *Banksia integrifolia* | 0.70 | 0.04 | 3 |
| *Plectranthus argenteatus* | 0.70 | 0.05 | 3 |
| *Cotyledon [orbiculata]* | 0.70 | 0.05 | 3 |
| *Cheilanthes fendleri* | 0.70 | 0.04 | 3 |
| *Nolina nelsonii* | 0.71 | 0.03 | 3 |
| *Astrolepis windhamii* | 0.71 | 0.04 | 6 |
| *Pleopeltis guttata* | 0.71 | 0.03 | 3 |
| *Butia eriostachya* | 0.71 | 0.02 | 3 |
| *Nerine undulata* | 0.72 | 0.02 | 3 |
| *Grevillea lanigera* | 0.73 | 0.04 | 3 |
| *Salvia muelleri* | 0.73 | 0.05 | 6 |
| *Sobralia fimbriata* | 0.73 | 0.03 | 3 |
| *Heuchera amoena* | 0.74 | 0.05 | 3 |
| *Peperomia obtusifolia* | 0.74 | 0.01 | 3 |
| *Paraceterach marantae* | 0.74 | 0.02 | 3 |
| *Clusia sp.* | 0.74 | 0.04 | 6 |
| *Selaginella peruviana* | 0.74 | 0.02 | 3 |
| *Platyterium bifurcatum* | 0.74 | 0.01 | 3 |
| *Anthurium watermalense* | 0.74 | 0.01 | 6 |
| *Selaginella pallescens* | 0.74 | 0.02 | 6 |
| *Blechnum caudatum* | 0.74 | 0.01 | 3 |
| *Actinopteris semiflabellata* | 0.74 | 0.03 | 3 |
| *Cavendishia lebroniae* | 0.74 | 0.04 | 3 |
| *Doryopteris elegans* | 0.74 | 0.00 | 3 |
| *Selaginella involvens* | 0.74 | 0.02 | 3 |
| *Calathea zebrina* | 0.75 | 0.01 | 3 |
| *Callistemon viminialis* | 0.75 | 0.01 | 3 |
| *Begonia venosa* | 0.75 | 0.02 | 3 |
| *Asplenium* | 0.75 | 0.02 | 3 |
| *Xanthorrhoea glauca* | 0.75 | 0.05 | 3 |
| *Eucalyptus forestiana* | 0.75 | 0.04 | 6 |
| *Comarostaphylis discolor* | 0.75 | 0.06 | 6 |
| *Cheilanthes bonariensis* | 0.76 | 0.02 | 6 |
| *Diplazium esculentum* | 0.76 | 0.02 | 3 |
| *Selaginella willendovii* | 0.76 | 0.01 | 3 |
| *Biophytum* | 0.76 | 0.00 | 3 |
| *Piper nigrum* | 0.76 | 0.01 | 3 |
| *Spathiphyllum floribundum* | 0.76 | 0.00 | 3 |
| *Selaginella unicata* | 0.76 | 0.02 | 3 |
| *Oatea acuminata* | 0.76 | 0.01 | 3 |
| *Kalanchoe pumila* | 0.76 | 0.01 | 6 |
| *Psilotum X intermedium* | 0.76 | 0.01 | 3 |
| *Adiantum macrophyllum* | 0.77 | 0.01 | 3 |
| *Coffea arabica* | 0.78 | 0.01 | 3 |
| *Anthurium brownii* | 0.78 | 0.00 | 3 |
| *Rhododendron arboreum* var. *arboreum* | 0.79 | 0.01 | 3 |
| *Acrostichum danæifolium* | 0.79 | 0.00 | 3 |
| *Yucca rostrata* | 0.80 | 0.00 | 3 |
| *Pteris altissima* | 0.81 | 0.00 | 3 |

**Table S2:** F_v_/F_m_ and n for each species sampled for fern diversity NPQ analysis.

| **Species** | **Mean $F_v/F_m$** | **SD** | **n** |
| --- | --- | --- | --- |
| *Pellaea rotundifolia* | 0.86 | 0 | 3 |
| *Athyrium filix-femina* var. *cyclosorum* | 0.83 | 0 | 3 |
| *Pleopeltis myriolepis* | 0.83 | 0.05 | 6 |
| *Acrostichum danaeifolium* | 0.82 | 0.05 | 6 |
| *Cyathea medullaris* | 0.82 | 0.06 | 6 |
| *Alsophila firma* | 0.81 | 0.03 | 6 |
| *Polystichum munitum* | 0.8 | 0.07 | 6 |
| *Woodwardia fimbriata* | 0.8 | 0.06 | 6 |
| *Osmunda spectabilis* | 0.79 | 0.07 | 6 |
| *Pleopeltis guttata* | 0.79 | 0.06 | 6 |
| *Todea barbara* | 0.79 | 0.06 | 6 |
| *Asplenium scolopendrium* | 0.78 | 0.08 | 6 |
| *Astrolepis laevis* | 0.78 | 0.08 | 6 |
| ***Astrolepis windhamii*** | **0.78** | **0.07** | **6** |
| *Astrolepis sinuata* | 0.78 | 0.06 | 6 |
| *Blechnum spicant* | 0.78 | 0.08 | 6 |
| ***Nicotiana benthamiana*** | 0.78 | 0.08 | 6 |
| *Actiniopteris semiflabellata* | 0.77 | 0.08 | 6 |
| *Adiantum capillus-veneris* | 0.77 | 0.07 | 6 |
| *Tectaria moorei* | 0.77 | 0.06 | 6 |
| *Diplazium esculentum* | 0.74 | 0.06 | 6 |
| *Adiantum aleuticum* | 0.73 | 0.09 | 6 |
| *Vandenboschia radicans* | 0.7 | 0.12 | 6 |
| *Salvinia auriculata* | 0.66 | 0.11 | 6 |

**Table S3:**
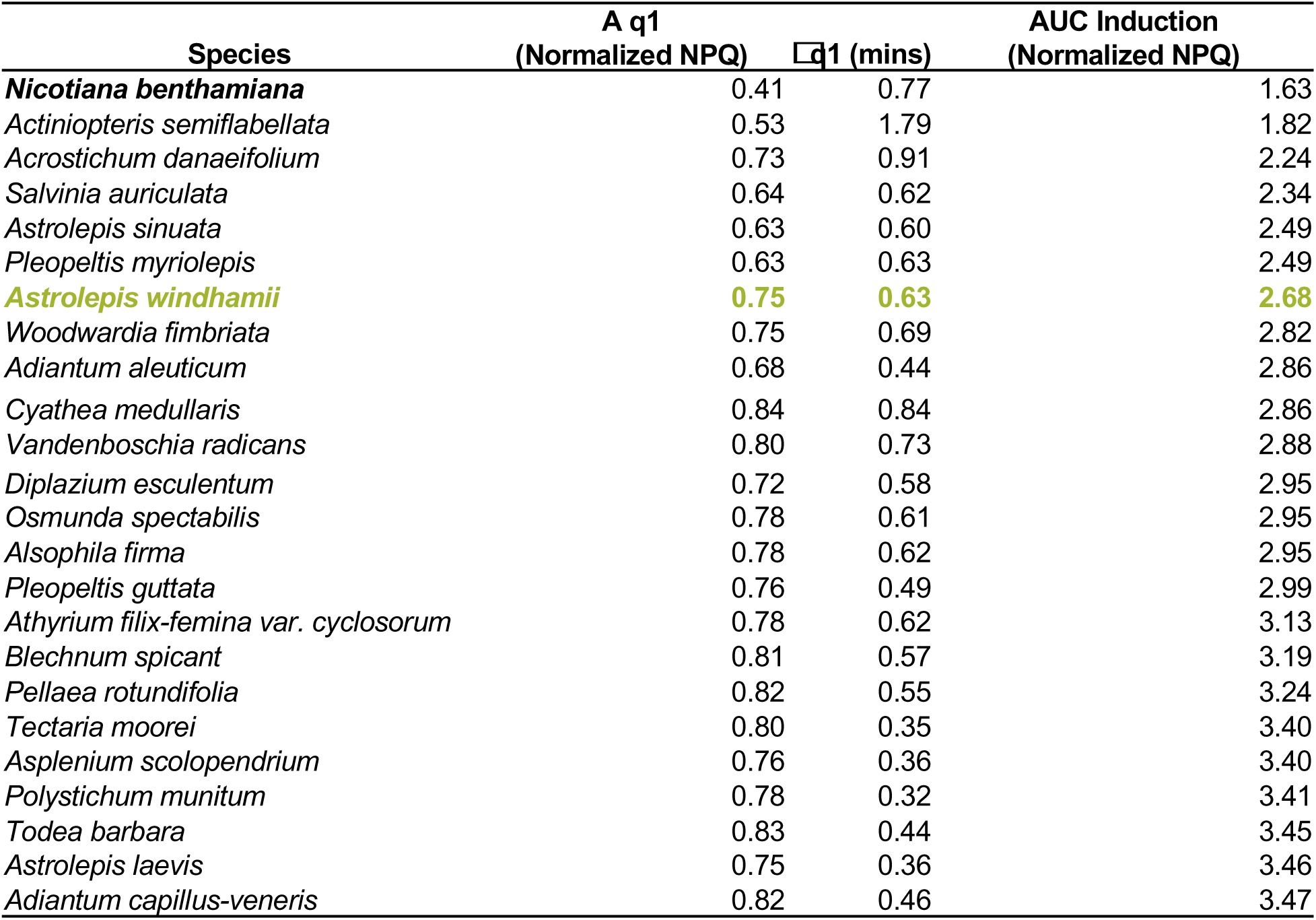
Exponential one-phase association curve for NPQ induction in fern diversity panel. 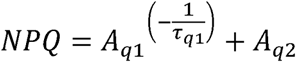

**Table S4:** Exponential decay best fit parameters for normalized NPQ relaxation (Period 1) in fern diversity panel. 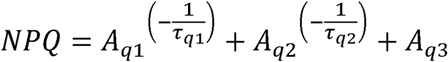

| Species | A q1 (Normalized NPQ) | $\tau_{q1}$ (mins) | A q2 (Normalized NPQ) | $\tau_{q2}$ (mins) | A q3 (Normalized NPQ) | AUC Dark (NPQ) |
| --- | --- | --- | --- | --- | --- | --- |
| *Salvinia auriculata* | 0.53 | 0.63 |  |  | 0.47 | 8.29 |
| *Pellaea rotundifolia* | 0.37 | 0.60 | 0.40 | 4.13 | 0.24 | 6.99 |
| *Adiantum aleuticum* | 0.63 | 0.39 | 0.07 | 2.86 | 0.30 | 6.38 |
| *Vandenboschia radicans* | 0.64 | 0.39 | 0.16 | 13.01 | 0.20 | 6.27 |
| ***Nicotiana benthamiana*** | **0.54** | **0.31** | **0.20** | **3.18** | **0.26** | **6.19** |
| *Actiniopteris semifabellata* | 0.77 | 0.48 |  |  | 0.22 | 5.06 |
| *Diplazium esculentum* | 0.56 | 0.35 | 0.27 | 2.28 | 0.17 | 4.72 |
| *Athyrium filix-femina* | 0.67 | 0.29 | 0.10 | 1.19 | 0.23 | 4.56 |
| *Tectaria moorei* | 0.80 | 0.32 |  |  | 0.20 | 4.53 |
| *Adiantum capillus-veneris* | 0.80 | 0.39 |  |  | 0.20 | 4.53 |
| *Astrolepis sinuata* | 0.72 | 0.34 | 0.14 | 3.91 | 0.13 | 4.37 |
| *Acrostichum danaeifolium* | 0.72 | 0.31 | 0.13 | 2.76 | 0.15 | 4.32 |
| *Todea barbara* | 0.82 | 0.36 |  |  | 0.18 | 4.21 |
| *Cyathea medullaris* | 0.75 | 0.43 | 0.13 | 4.18 | 0.13 | 4.21 |
| *Blechnum spicant* | 0.75 | 0.45 | 0.13 | 3.83 | 0.13 | 4.13 |
| *Osmunda spectabilis* | 0.75 | 0.39 | 0.11 | 5.91 | 0.14 | 4.12 |
| *Pleopeltis myriolepis* | 0.69 | 0.40 | 0.18 | 2.01 | 0.14 | 4.11 |
| *Alsophila firma* | 0.60 | 0.27 | 0.27 | 1.55 | 0.13 | 3.93 |
| *Pleopeltis guttata* | 0.80 | 0.47 | 0.08 | 2.80 | 0.13 | 3.70 |
| *Astrolepis laevis* | 0.79 | 0.27 | 0.09 | 5.81 | 0.12 | 3.70 |
| *Asplenium scolopendrium* | 0.85 | 0.24 |  |  | 0.15 | 3.62 |
| *Woodwardia fimbriata* | 0.81 | 0.29 | 0.07 | 3.74 | 0.12 | 3.50 |
| *Polystichum munitum* | 0.86 | 0.34 |  |  | 0.14 | 3.36 |
| ***Astrolepis windhamii*** | **0.77** | **0.30** | **0.12** | **2.52** | **0.11** | **3.25** |

**Table S5:** Exponential decay best fit parameters for normalized NPQ relaxation (Period 2) in fern diversity panel. 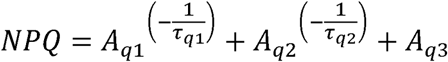

| Species | A q1<br>(Normalized NPQ) | τ q1 (mins) | A q2<br>(Normalized NPQ) | τ q2 (mins) | A q3<br>(Normalized NPQ) | AUC Dark (NPQ) |
| --- | --- | --- | --- | --- | --- | --- |
| *Salvinia auriculata* | 0.53 | 0.63 |  |  | 0.66 | 8.79 |
| *Adiantum aleuticum* | 0.74 | 0.48 |  |  | 0.51 | 7.87 |
| *Pellaea rotundifolia* | 0.59 | 1.12 |  |  | 0.48 | 7.41 |
| ***Nicotiana benthamiana*** | **0.73** | **0.67** |  |  | **0.44** | **6.93** |
| *Vandenboschia radicans* | 0.68 | 0.51 |  |  | 0.41 | 6.52 |
| *Athyrium filix-femina* var. *cyclosorum* | 0.53 | 0.64 | 0.41 | 0.19 | 0.39 | 5.83 |
| *Actiniopteris semiflabellata* | 0.77 | 0.39 |  |  | 0.28 | 5.15 |
| *Adiantum capillus-veneris* | 0.80 | 0.39 |  |  | 0.30 | 5.01 |
| *Pleopeltis myriolepis* | 0.90 | 0.79 |  |  | 0.19 | 4.96 |
| *Tectaria moorei* | 0.88 | 0.28 |  |  | 0.28 | 4.91 |
| *Diplazium esculentum* | 0.60 | 0.66 | 0.24 | 0.15 | 0.25 | 4.82 |
| *Blechnum spicant* | 0.85 | 0.50 |  |  | 0.27 | 4.81 |
| *Todea barbara* | 0.84 | 0.32 |  |  | 0.26 | 4.47 |
| *Acrostichum danaeifolium* | 0.50 | 0.61 | 0.29 | 0.16 | 0.24 | 4.41 |
| *Cyathea medullaris* | 0.86 | 0.47 |  |  | 0.21 | 4.36 |
| *Astrolepis sinuata* | 0.53 | 0.58 | 0.27 | 0.13 | 0.23 | 4.26 |
| *Osmunda spectabilis* | 0.84 | 0.38 |  |  | 0.24 | 4.19 |
| *Pleopeltis guttata* | 0.89 | 0.59 |  |  | 0.19 | 4.17 |
| *Asplenium scolopendrium* | 0.89 | 0.30 |  |  | 0.22 | 4.13 |
| *Alsophila firma* | 0.50 | 0.69 | 0.34 | 0.17 | 0.20 | 4.08 |
| *Woodwardia fimbriata* | 0.90 | 0.31 |  |  | 0.21 | 3.98 |
| *Polystichum munitum* | 0.89 | 0.34 |  |  | 0.21 | 3.84 |
| *Astrolepis laevis* | 0.83 | 0.29 |  |  | 0.21 | 3.80 |
| ***Astrolepis windhamii*** | **0.87** | **0.35** |  |  | **0.18** | **3.29** |

**Fig. S1:**
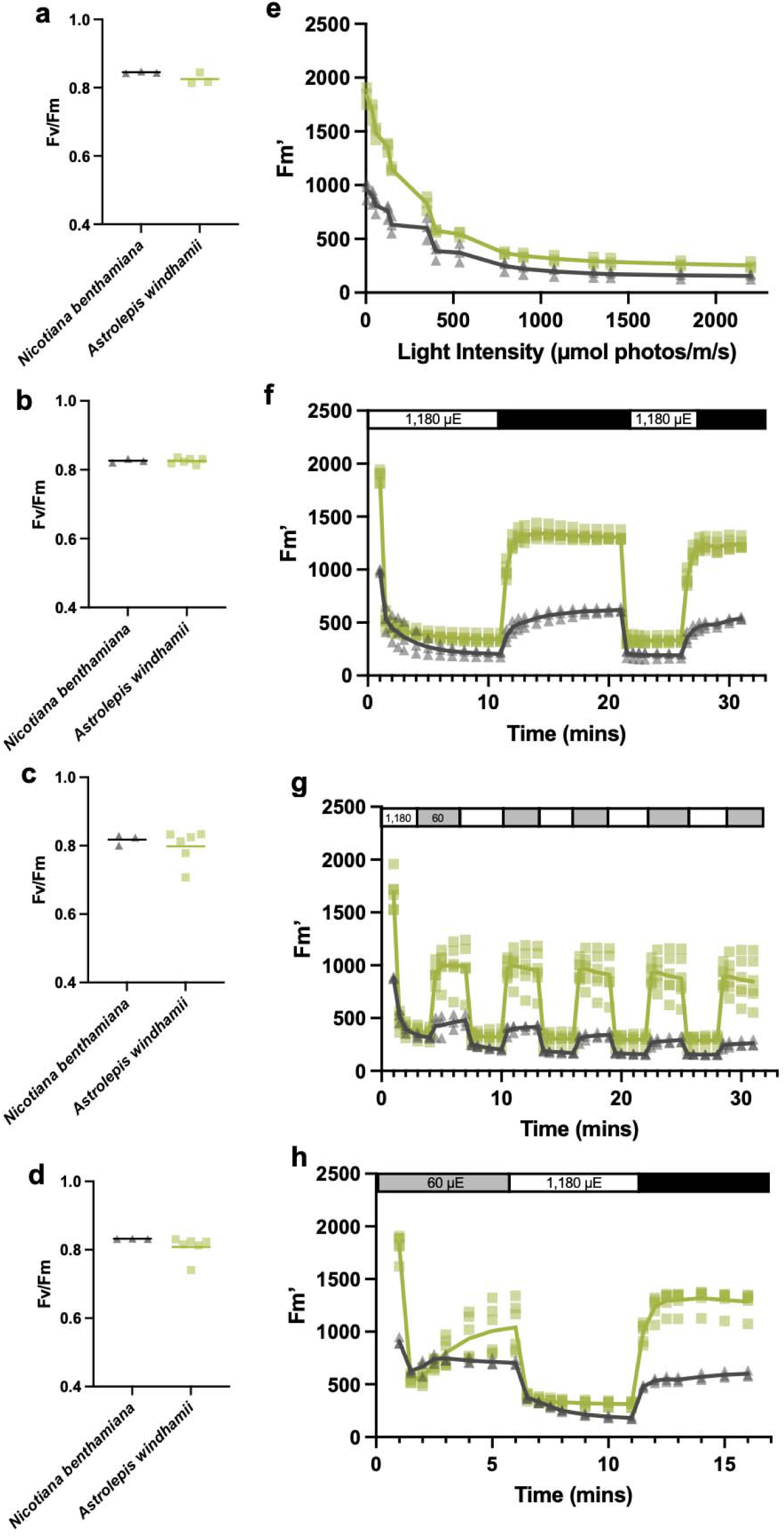
Astrolepis NPQ analysis, F_v_/F_m_ and F_m_’ data. **a.** F_v_/F_m_ for light response curve data (Fig 3) in Astrolepis and tobacco (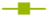 Astrolepis (n=3), 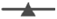 tobacco (n=3)). **b.** Raw F_m_’ for NPQ data (Fig 3a**-b**) in Astrolepis. **c.** F_v_/F_m_ for NPQ data (Fig 4a**-b**) in Astrolepis and tobacco (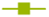 Astrolepis (n=6), 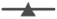 tobacco (n=3)). **d.** Raw F_m_’ for NPQ data (Fig 4a**-b**) in Astrolepis and tobacco (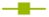 Astrolepis (n=6), 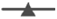 tobacco (n=3)). **e.** F_v_/F_m_ for NPQ data (Fig 4e**-f**) in Astrolepis and tobacco (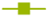 Astrolepis (n=6), 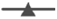 tobacco (n=3)). **f.** Raw F_m_’ for NPQ data (Fig 4e**-f**) in Astrolepis and tobacco (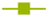 Astrolepis (n=6), 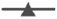 tobacco (n=3)). **g.** F_v_/F_m_ for NPQ data (Fig 4i**-j**) in Astrolepis and tobacco (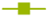 Astrolepis (n=6), 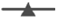 tobacco (n=3)). **h.** Raw F_m_’ for NPQ data (Fig 4i**-j**) in Astrolepis and tobacco (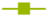 Astrolepis (n=6), 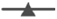 tobacco (n=3)).

**Fig. S2:**
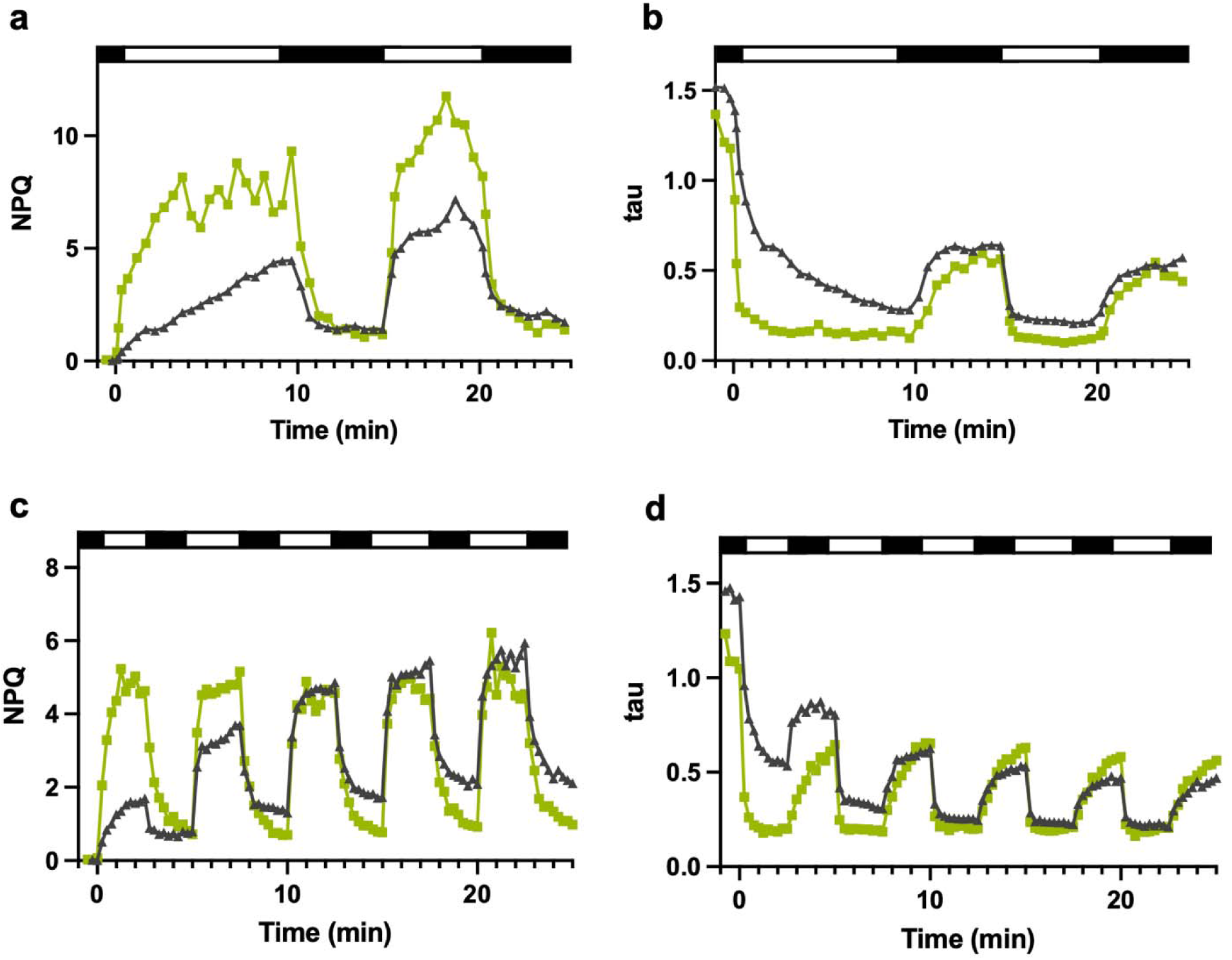
Time Correlated Single Photon Counting (TCSPC) data for Astrolepis. **a.** Mean NPQ calculated from TCSPC (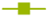 Astrolepis (n=4), 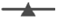 tobacco (n=3)) during a 10 min HL regime (1,400 µmol photons m^-2^ s^-1^) followed by 10 min darkness, followed by a 5 min HL-dark cycle. **b.** Mean tau (fluorescence lifetime) in **a**. **c.** Mean NPQ calculated from TCSPC (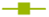 Astrolepis (n=4), 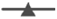 tobacco (n=3)) during 5 cycles of a 2.5 min HL regime (1,400 µmol photons m^-2^ s^-1^) followed by 2.5 min darkness. **d.** Mean tau (fluorescence lifetime) in **c**.

